# Diverse Hotspot Thermal Profiling Methods Detect Phosphorylation-Dependent Changes in Protein Stability

**DOI:** 10.1101/2021.05.01.441686

**Authors:** Benjamin D. Stein, Jun X. Huang, David Wu, Lewis C. Cantley, Raymond E. Moellering

## Abstract

Hotspot thermal profiling (HTP) methods utilize modified-peptide level information in order to interrogate proteoform-specific stability inside of live cells. The first demonstration of HTP involved the integration of phosphopeptide enrichment into a TMT-based, single-LC separation thermal profiling workflow^1^. Here we present a new ‘label-fractionate-enrich’ (LFE)-HTP method that involves high-pH reverse phase fractionation of TMT-labeled peptides prior to phosphopeptide enrichment, followed by peptide detection and quantitation using multi-notch LC-MS^3^. We find that LFE-HTP, while more resource intensive, improves the depth and precision of (phospho)proteoform coverage relative to the initial published HTP workflow. The fraction of detected phosphorylation sites that are significantly perturbed in this new dataset are consistent with those seen in our previous study, as well as those published by others, when compared head-to-head with the same analysis pipelines. Likewise, many ‘hotspot’ phosphorylation sites identified in our paper are consistently reproduced by LFE-HTP as well as other modified HTP methods. The LFE-HTP dataset contains many novel ‘hotspot’ phosphorylation sites that regulate the stability of diverse proteins, including phosphosites in the central glycolytic enzyme Aldolase A that are associated with monomer-to-oligomer formation, enzymatic activity and metabolic regulation in cancer cells. Our comparative analyses confirm that several variants of the HTP method can track modified proteoforms in live cells to detect and prioritize PTM-dependent changes in protein stability that may be associated with function.

## Introduction

Thermal profiling approaches permit massively parallel biophysical measurements to be made in whole proteome (i.e. lysates), live cells and tissue samples^2-4^. These methods utilize pulsed proteome denaturation by exposing aliquots of cells to variable temperatures, followed by fast freeze-thaw, centrifugation of insoluble proteins, and isolation of the soluble supernatant for mass spectrometric analysis. In almost all cases, a form of isotopic barcoding is employed to make quantitative measurements of protein abundance in samples exposed to different temperatures; this peptide-level information is ultimately used to calculate protein half-maximal melting temperatures (T_m_)^5^ or solubility indices^6^. While all measurements are made at the peptide level, the vast majority of thermal profiling studies collapse peptide-level information to yield protein T_m_ values and their differences upon exposure to perturbations like small molecule ligands, metabolites, or other stimuli^7-10^. Recently, we reported a method, hotspot thermal profiling (HTP), which captures modified-peptide level information in order to specifically interrogate the biophysical and biochemical properties of unique proteoforms in cells^1^.

Our first report of the HTP method was applied to protein phosphorylation, and yielded multiple conclusions, including: 1) Modified-peptide level thermal profiling information allows for biophysical interrogation of proteoforms in live cells; 2) Site-specific posttranslational modifications (PTMs) can significantly affect proteoform T_m_ relative to the bulk-unmodified pool of the same protein; 3) These “hotspot” sites may represent functionally relevant PTMs that

perturb protein structure and function; 4) Uncharacterized modifications that impact inter- and intramolecular interactions can be discovered and prioritized for subsequent functional studies. Since the initial publication of this method, other reports have used similar thermal profiling approaches to show that redox-mediated modifications, protein phosphorylation and site-specific mutations can affect proteoform stability relative to the ‘parent’ or unmodified pool of given protein^11-14^. Notably, the defined pools of proteoforms that are compared differ among reports in the literature, and therefore the biochemical and statistical comparisons are equally varied. In the HTP method, we deliberately compare specific phosphomodiforms (defined by unequivocal presence of a phosphorylation site in detected phosphopeptides) relative to the unmodified bulk parent protein pool detected in cells (defined as bulk, unmodified). This measurement therefore compares the thermal stability of a subset of proteoforms that contain that modification site to the bulk protein population, which is also an inclusive set of proteoforms (Suppl. Fig. 1). Another possibility is to perform a one-to-one comparison of the modified and unmodified versions of a protein using the same peptide regions as markers for those respective protein pools. Finally, others have analyzed these datasets by comparing each modified peptide and every other detected peptide within a protein, one-by-one (i.e. a single peptide vs. dozens or hundreds of peptides from the same protein)^15^. While the varied workflows and data analysis pipelines seem similar, they entail different biochemical comparisons, and likewise could require different statistical considerations to account for multiple-hypothesis testing. We were interested in testing how these factors affect method performance, and therefore developed new biochemical workflows and compared different data analysis pipelines. Herein, we present a new ‘label-fractionate-enrich’ version of the HTP method that improves depth of (phospho)proteome coverage and measurement precision. We compare the global trends and specific ‘hotspot’ phosphorylation site-dependent measurements to our previously published dataset, as well as datasets that have been reported by other groups. Finally, we consider the impact of different data analysis pipelines on the interpretation of HTP-like datasets and the inferred prevalence of phosphorylation sites that significantly impact protein stability.

## Results

Since the first report of the HTP method, we have explored several modifications to the HTP workflow that could affect the depth of (phospho)proteome coverage, precision in sample processing, LC-MS measurement precision, and data interpretation. For example, we hypothesized that separate TMT-labeling of bulk and phosphopeptide fractions, use of TiO_2_ spin-tips and LC-MS/MS acquisition with a single chromatographic separation could collectively contribute to lower measurement precision in our original workflow. To modify these factors and compare to our original method, we developed a new ‘label-fractionate-enrich’ (LFE-HTP) workflow (Fig. 1A). This workflow involves TMT-labeling and pooling of all peptides from different temperature fractions, high-pH reverse-phase fractionation, and subsequent splitting of each fraction for bulk protein analysis (5%) and TiO_2_ enrichment (95% of each fraction).

**Figure 1:**
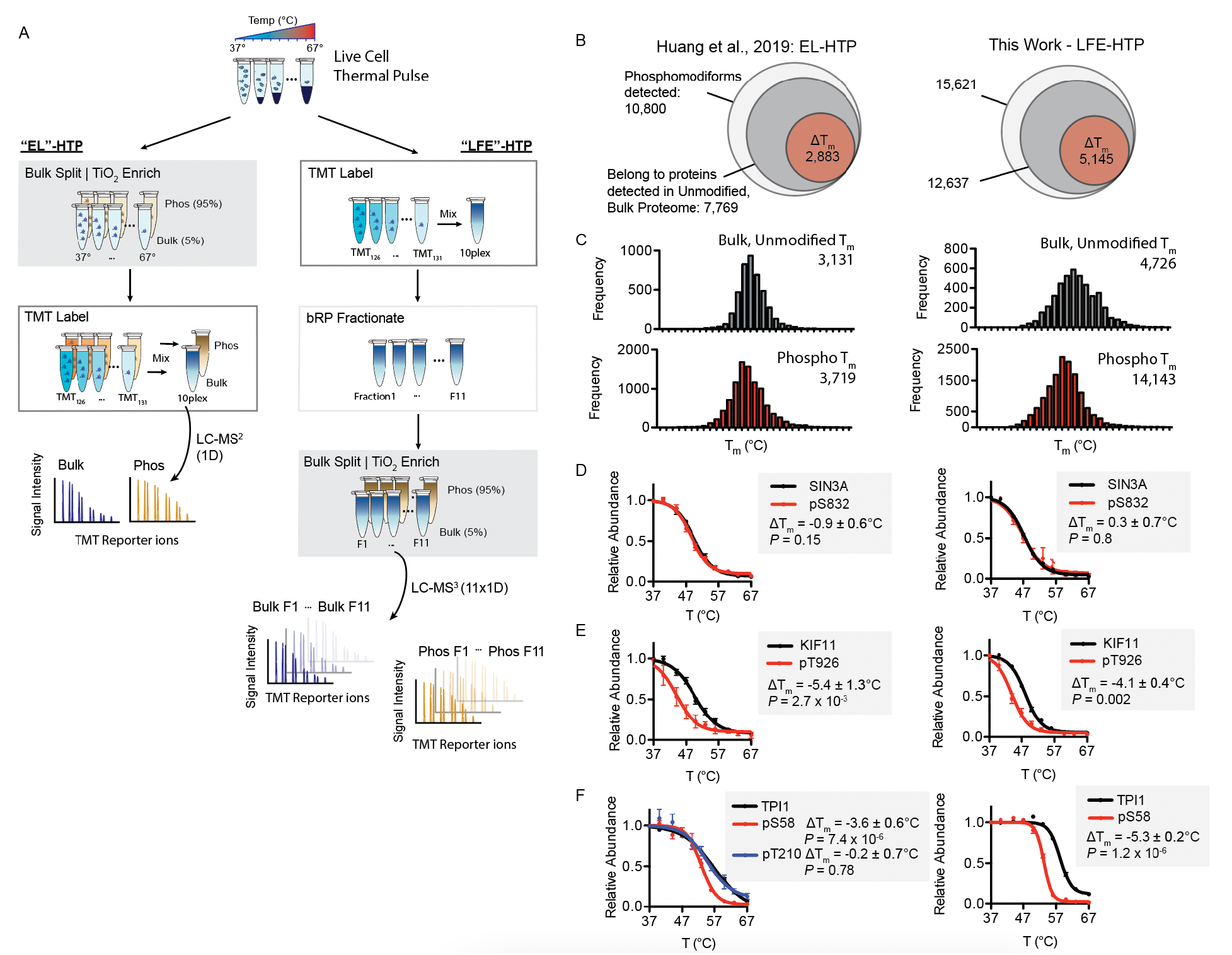
Comparison of variant HTP workflows. **A)** Proteomic workflow schematics of two HTP variants: EL (enrich-label) and LFE (label-fractionate-enrich). **B**) Total number of unique phosphosites and resulting ΔT_m_ values detected in each HTP dataset. **C**) Histograms depicting global distribution of unmodified, bulk protein and phosphomodiform T_m_ values in each HTP dataset. **D-F**) Representative T_m_ curves of bulk, unmodified protein (black) and indicated phosphomodiforms (red) from SIN3A (D), KIF11 (E), and TPI1 (F). Only curves with *R*^2^ > 0.8 are plotted. Curves and error bars in **D-F** correspond to T_m_ mean and s.e.m.; ΔT_m_ *P*-values are calculated with two-sided *t*-test.

Therefore, each LFE-HTP dataset is derived from 22 LC-MS/MS runs and aided by the use of multi-notch MS^3^ acquisition to reduce ratio-compression and improve precision in TMT quantification of phosphopeptides^16, 17^.

Among four LFE-HTP biological replicates prepared with live HEK293T cells, we measured 4,726 ‘bulk, unmodified’ protein T_m_ values, and 14,143 phosphosite-specific proteoform (phosphomodiform) T_m_ values, which represented a significant increase in both the bulk, unmodified and phosphosite identifications and T_m_ measurements relative to our published “enrich-then-label” (EL)-HTP dataset (Fig. 1B-C). Importantly, this dataset was derived from ∼10x the number of LC-MS runs, and was performed on a different instrument with different acquisition settings. The global relationships previously observed between phosphosite environment and proteoform stability in the LFE-HTP dataset were similar to those reported in our original manuscript with the EL-HTP method (Suppl. Fig. 2). As predicted, a single labeling event of the unenriched proteome with increased TMT reagent use, orthogonal fractionation (22 vs. 2 MS runs per sample) prior to phospho-enrichment and multi-notch MS^3^ analysis resulted in higher technical and biological precision in LFE-HTP measurements (Suppl. Fig. 3). Notably, the increased precision and deeper phosphoproteome coverage of the LFE-HTP dataset did not correspond with a lower fraction of phosphorylation sites that significantly perturbed proteoform stability relative to the bulk, unmodified pool of the same protein using our published data analysis pipeline. On the contrary, ∼26% of phosphosites in our LFE-HTP dataset have a ΔT_m_ > 1.5°C and a *p-*value < 0.05 – very similar to that found in our original dataset and in datasets published by others (*vide infra*).

Within this dataset we observed replication of many ‘hotspot’ phosphosite ΔT_m_ values previously measured and reported in the EL-HTP dataset. pS832 in SIN3A, which did not affect proteoform stability in our original dataset, likewise showed overlaid curves in the more precise LFE-HTP dataset (Fig. 1D). Other phosphosites highlighted in our first dataset, including pT926 in KIF11 and pS58 in TPI1, show highly similar and statistically significant ΔT_m_ values across HTP workflows and datasets (Fig. 1E-F). We also observed similar changes in stability for several phosphosites in the N-terminal tail of 4EBP1 as reported in our previous publication (Suppl. Fig. 4). Intriguingly, the ‘bulk, unmodified’ protein pool and several of its modified proteoforms have a clear and consistent unfolding curve that plateaus in the ∼30-50% abundance range relative to aliquots exposed to low temperatures. We note this protein in particular as the curve fitting assumptions and requirements imposed by some analysis pipelines cannot (and do not) capture this protein or these sites, as they do not completely “melt.” The subjective decision to include or exclude such proteins and sites could have a significant effect on perceived trends within datasets.

The LFE-HTP dataset contains many novel phosphorylation sites that significantly impact the protein thermal stability of proteins in diverse ways (Supplementary Tables 1-2). As previously presented, hotspot sites in this dataset likely impact proteoform stability by altering intramolecular interactions, protein complex interfaces, protein-metabolite interactions, and combinations thereof. For example, the LFE-HTP dataset identified pS46, pS360, and pY364 belonging to Aldolase A (ALDOA), each of which significantly decreased the melting point of their corresponding phosphomodiforms, despite being far from each other in primary and tertiary structure (Fig. 2A). Serine 46 is located on the same interface as the active site pocket, and is adjacent to Arg43, which has been shown to be involved in ALDOA recruitment to F-actin and ensuing regulation of enzyme activity and metabolism^18^ (Fig. 2B). Phosphorylated S360 and Y364 are located in the C-terminal tail of ALDOA, which has been shown to adopt a significantly altered conformation in monomeric and tetrameric forms of the enzyme. In the monomer, this peptide stretch is bound on the surface of the enzyme, with Y364 buried into the active site^19^ (Fig. 2B). In the tetrameric form, which is associated with decreased activity^20^, both S360 and Y364 are located away from the protein surface and the surrounding C-terminal peptide is involved in the dimer interface (Fig. 2C). Intriguingly, no electron density is present for the residues surrounding S46 and R43 in the tetramer structure, despite their proximity to the active site. While no specific function has been reported for these sites in ALDOA, the strong destabilization of these modified proteoforms indicates that they may disrupt active site architecture, quaternary structure, protein-protein interactions or combinations thereof, that regulate protein abundance, activity and localization. The unbiased identification of potential hotspot regulatory sites like these is an advantage of the HTP approach, and more detailed interrogation of these and other sites in this dataset is warranted.

**Figure 2:**
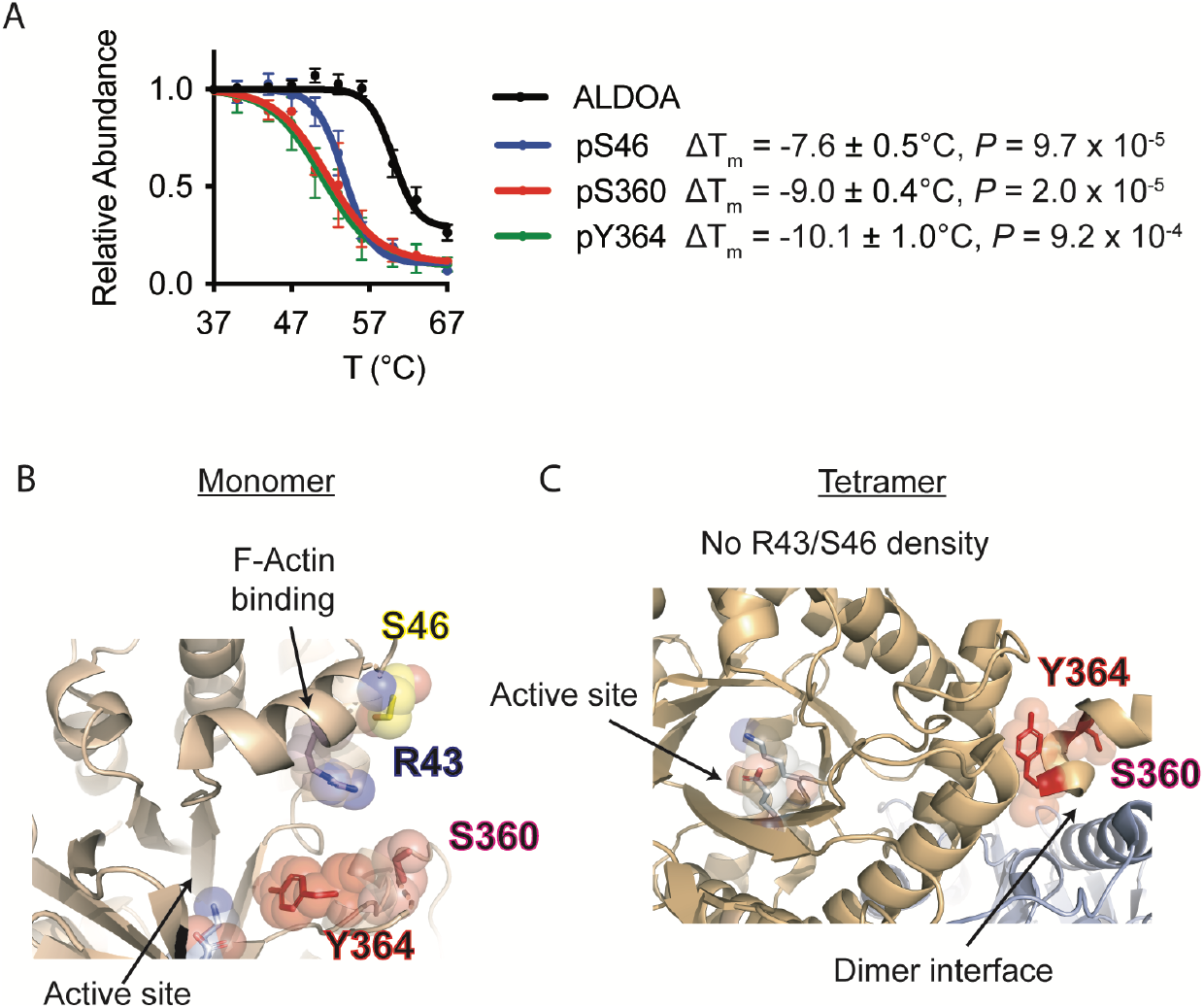
LFE-HTP identifies phosphosites with significant thermal shifts in aldolase. **A)** Phosphosites pS46, pS360, and pY364 induce significant and similar negative thermal stability shifts on aldolase A. **B)** In the monomeric conformation (PDB: 1ALD), S46 (yellow) is adjacent to R43 (blue) which has been shown to be involved in ALDOA recruitment to F-actin, meanwhile, C-terminal Y364 (red) and S360 (pink) are packed inside the enzyme active site. **C)** In the tetramer structure Y364 (red) and S360 (pink) are far away from the active site (blue) but packed next to the dimer interface (PDB: 5KY6). Curves and error bars in **A)** correspond to T_m_ mean and SEM; Only curves with *R*^2^ > 0.8 are plotted. ΔT_m_ *P*-value is calculated using 2-sided student’s t-test.

Following our publication of the EL-HTP method, several other reports have applied the concept in various forms to measure the effect of protein modifications or mutations on protein stability. Among these, a method similar to our LFE-HTP approach was reported to employ up-front TMT labeling and fractionation after phosphopeptide enrichment, alongside other significant differences in the subsequent biochemical workflow and data analysis pipeline^15^.

Nonetheless, this approach and dataset provides an appropriate comparator to the EL-HTP and LFE-HTP methods. We were particularly intrigued by the conclusion that their methodology yielded substantially fewer statistically significant shifted phosphorylation sites (∼3%) relative to our original report in light of the fact that our analogous LFE-HTP dataset did not. To understand this discrepancy, we directly analyzed their dataset using our published analysis pipeline, which reproduced essentially identical T_m_ curves and ΔT_m_ values for modified proteoforms highlighted in their dataset (Fig. 3A). Therefore, despite differences in the overall analysis pipelines and statistical treatment of the ensuing data (i.e., comparing a phosphopeptide to an aggregate protein-level T_m_, or comparing a phosphopeptide to each unmodified peptide one-by-one), their analysis pipeline objectively reproduces the curves and shifts in thermal stability generated by our published analysis method, and vice versa.

**Figure 3:**
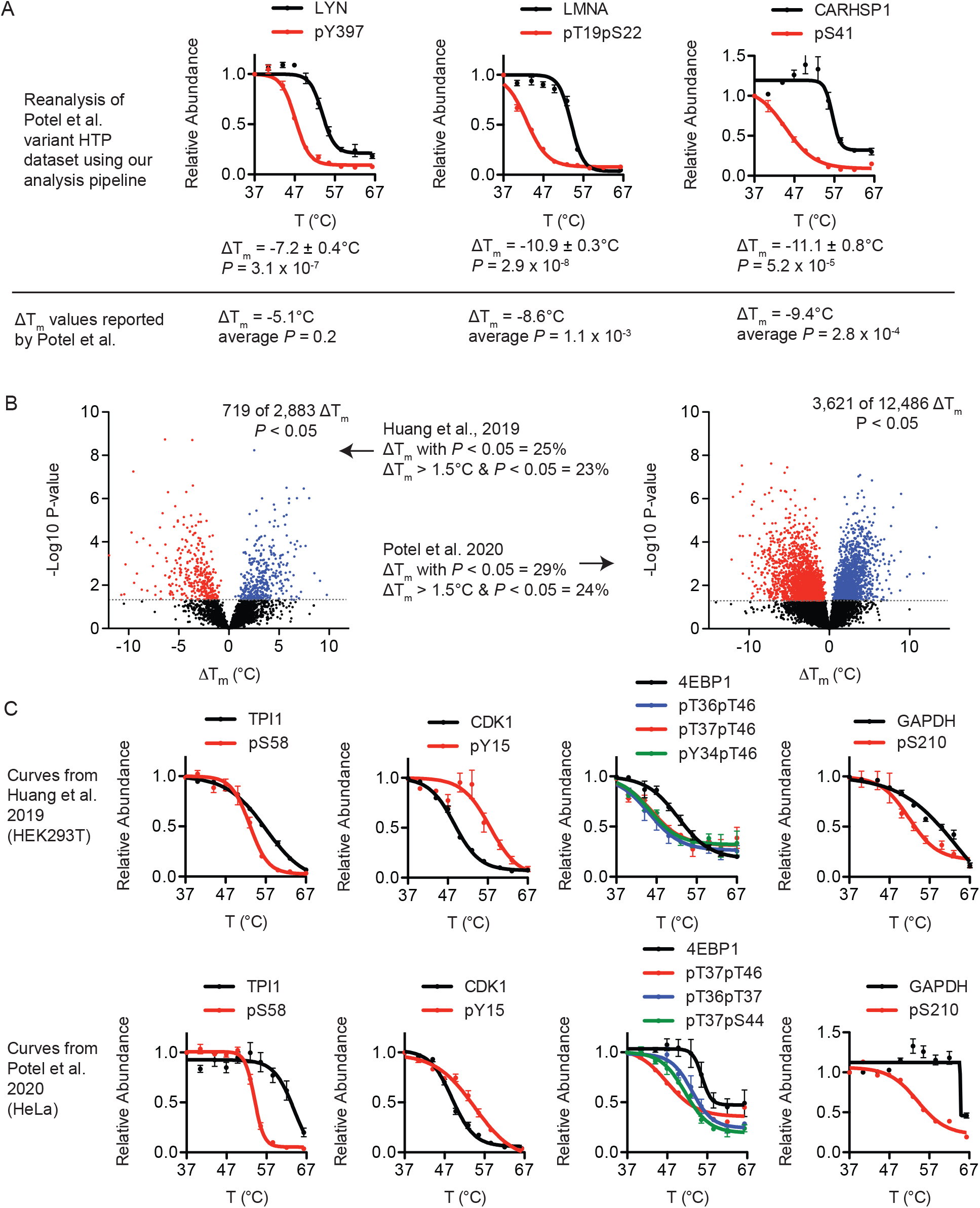
Compare data analysis pipelines. **A)** Reanalysis of dataset produced by Potel et al. with our published data analysis pipeline directly reproduced the melting curves highlighted in their dataset and yielded essentially identical ΔT_m_ values for specific reported modiforms. ΔT_m_ values in (A) are directly reported by Potel et al.^1^ for comparison. **B)** Global ΔT_m_ volcano plots from Huang et al., 2019 (left) and Potel et al. report (right). The Potel et al. dataset produces contains a near-identical (24%) percentage of significantly shifted phosphomodiforms relative to our published dataset (23%) when data are analyzed head-to-head with our published analysis pipeline. **C**) Representative melting curves previously reported using EL-HTP, including TPI1 (modiform pS58), CDK1 (modiform pY15), 4EBP1 (several modiforms in the N-terminal tail) and GADPH (pS210) were reproduced in the Potel et al. dataset.

Since both analysis pipelines produce essentially identical shifts on a site-by-site and protein-by-protein basis, we performed a global reanalysis of their data and compared to our published profile and newly generated LFE dataset. This objective reanalysis confirmed that their data harbors just as many significantly shifted phosphorylation sites as our original dataset. In particular, their alternative HTP workflow yields a dataset in which ∼24% of detected phosphorylation sites have a ΔT_m_ > 1.5°C and a *p-*value < 0.05 (Fig. 3B); this value was 23% in our original study, and 26% in our analogous LFE-HTP dataset. Diving deeper into their dataset, we also find that ΔT_m_’s for many of the anecdotal ‘hotspot’ phosphorylation sites detected in both our EL- and LFE-HTP datasets are reproduced in their dataset. While not presented in their analysis, their HTP workflow detects significant shifts in the stability of phosphomodiforms of TPI1 (pS58), CDK1 (pY15), 4EBP1 (several sites in the N-terminal tail) and GADPH (pS210; Fig. 3C). The last three are notable, as sites like pY15 in CDK1 are not captured by their search pipeline due to omission of sites that are completely conserved among protein isoforms, and the observed 4EBP1 and GAPDH shifts are not captured due to arbitrary cutoffs imposed on curve shape and plateaus. Taken together, these comparisons demonstrate that their conclusion of a more precise HTP method yields a vastly different fraction of shifted sites in the proteome – 3%, to be specific – is not objectively true. Instead, what drives this claim is their use of a divergent downstream computational processing pipeline. As expected, application of a more stringent statistical adjustment will bring the perceived significant hits down in their dataset and ours, but they come down uniformly (Suppl. Data Fig. 5). This comparison is useful as it is important to weigh the relative merits of including more or less stringent significance filters for hypothesis-generating experiments, and it is unsurprising that application of divergent significance comparisons to a dataset after the fact will yield different results.

## Discussion

Herein we presented a new hotspot thermal profiling method to measure the thermal stability of modified proteoforms in live cells. We confirmed that the application of increased fractionation, concomitant isotopic labeling reagents, instrument time and multi-notch MS^3^ quantification improves the depth of phosphoproteome coverage and precision of T_m_ measurements. The ensuing dataset reproduced many anecdotal hotspot sites from our first report, as well as in datasets produced by others. Furthermore, the LFE-HTP method identified 3 novel destabilizing sites in the central glycolytic enzyme ALDOA that we propose could disrupt the inactive tetramer or inhibitory interactions with F-actin providing mechanistic insight to altered enzymatic activity, which is further supported by previous studies. Our comparison of global trends suggest that a significant fraction of phosphorylation sites meaningfully perturb the

stability of their parent proteins. This notion is supported by the fact that two independent and more precise HTP methods show that approximately 25% of detected sites display a ΔT_m_ > 1.5°C and a *p-*value < 0.05 relative to the bulk, unmodified protein pool when analyzed with the same published analysis approach. Of course, one can impose different cutoffs for what could be considered as ‘significant’ and in doing so change the subjective interpretation of these datasets. However, since a primary purpose of HTP-like methods is to generate hypotheses for further study with global profiling (i.e. perturb the system and track proteoform-dependent effects) or through alternative methods, we posit that reporting directly on site-specific changes in stability alone is of primary importance. A prototypical example of this dichotomy is evident with the well-characterized 4EBP1 sites identified in our original manuscript, reproduced by our optimized LFE approach and observed in datasets now published by others. These sites are omitted using more stringent analysis pipelines, yet they are highly reproducible and represent validated regulatory events on this translational repressor protein. Moving forward, we expect that like other proteomic methods, future advances and derivations will improve upon those discussed here to enable continued application of hotspot thermal profiling to identify and characterize functionally important PTMs.

## Acknowledgements

We thank D. Huo and T. Harrison for discussions surrounding data analysis. Financial support for this work is from NCI P01CA120964, R35CA197588 (to L.C.C.) and R00CA175399 (to R.E.M.).

## Author contributions

B.D.S and R.E.M. conceived of the study. B.D.S supervised the study, designed, performed and analyzed experiments. J.X.H. performed cell-based and mass spectrometry experiments and analyzed data. D.W. performed cell-based and mass spectrometry experiments. L.C.C. oversaw part of the study. R.E.M. supervised the study, designed, analyzed experiments, and wrote the manuscript with input from all authors.

## Competing Interest Statement

The authors declare that they have no competing interests.

## Supplemental Materials

**Suppl. Fig. 1.**
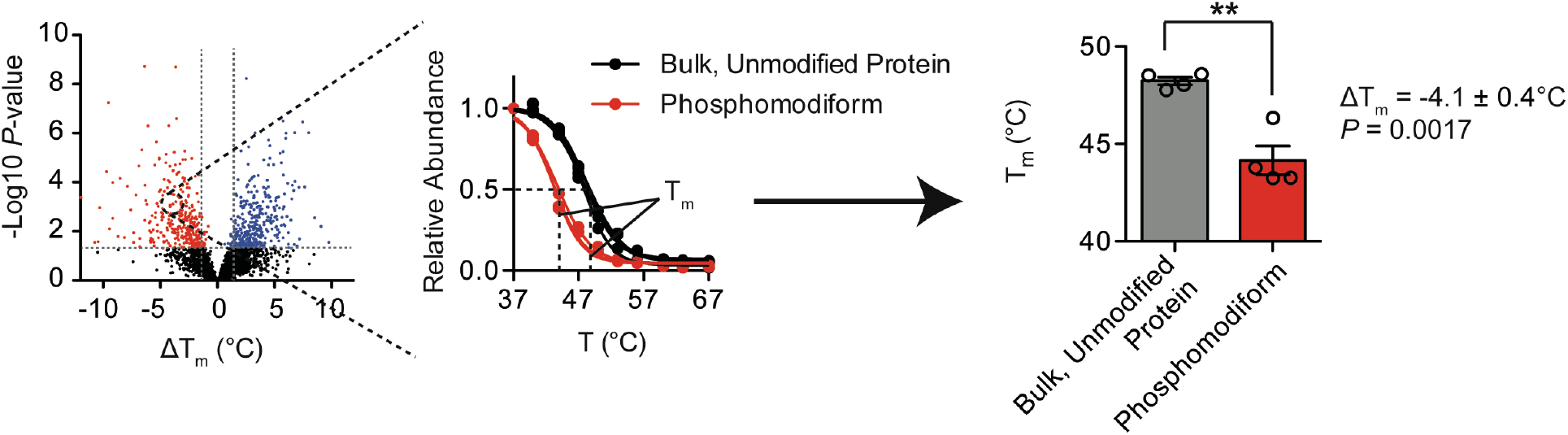
Schematic of statistical comparisons made to analyze HTP data. Schematic of statistical comparisons used which compares the means and error surrounding bulk T_m_ and phosphomodiform T_m_ and assesses whether a site significantly alters protein stability. A two-side t-test is used to compare the T_m_ values.

**Suppl. Fig. 2.**
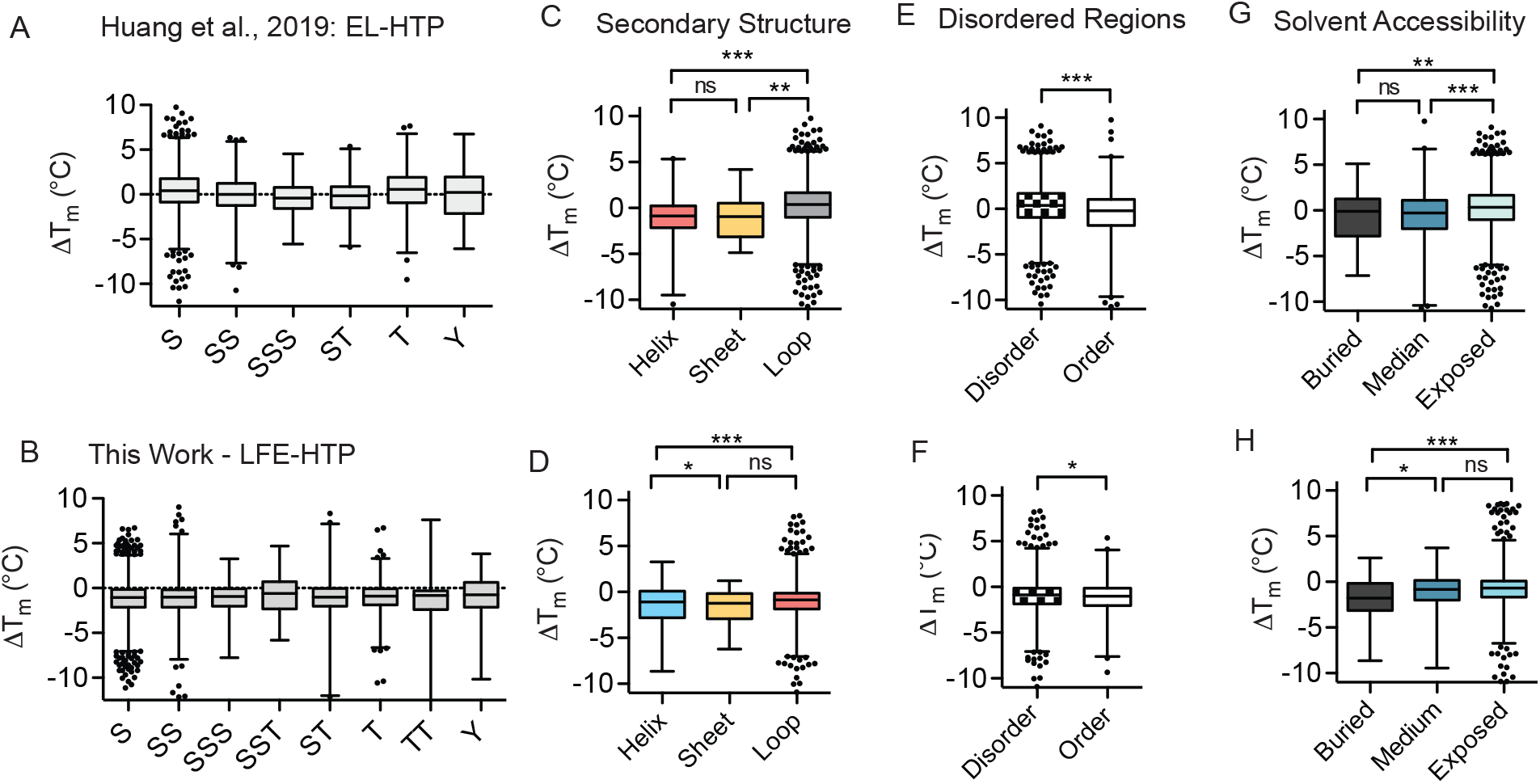
Comparing global relationships between local phosphosite environment and altered phosphomodiform stability of the two HTP variants. **A-B**) Distributions of ΔT_m_ values for tryptic peptides containing indicated phospho-amino acids or coincidental combinations thereof for Huang et al 2019 EL-HTP (A) and LFE-HTP (B). **C-H**) Comparisons of ΔT_m_ values and predicted secondary structure elements (C-D), ordered structural elements surrounding the phosphosite of interest (G-I) and solvent accessibility (E-F) of the three datasets respectively. For secondary structure (C-D), disordered regions (E-F) and solvent accessibility (G-H) analyses performed for LFE-HTP datasets, only proteins detected in the original Huang et al., 2019 publication are included in the analyses. For C-H), \*\*\**P* ≤ 0.0001, \*\**P* ≥ 0.001, \**P* ≥ 0.05, two-sided t-test. Box plots (median, 1–99%) are shown with outliers shown as data points.

**Suppl. Fig. 3.**
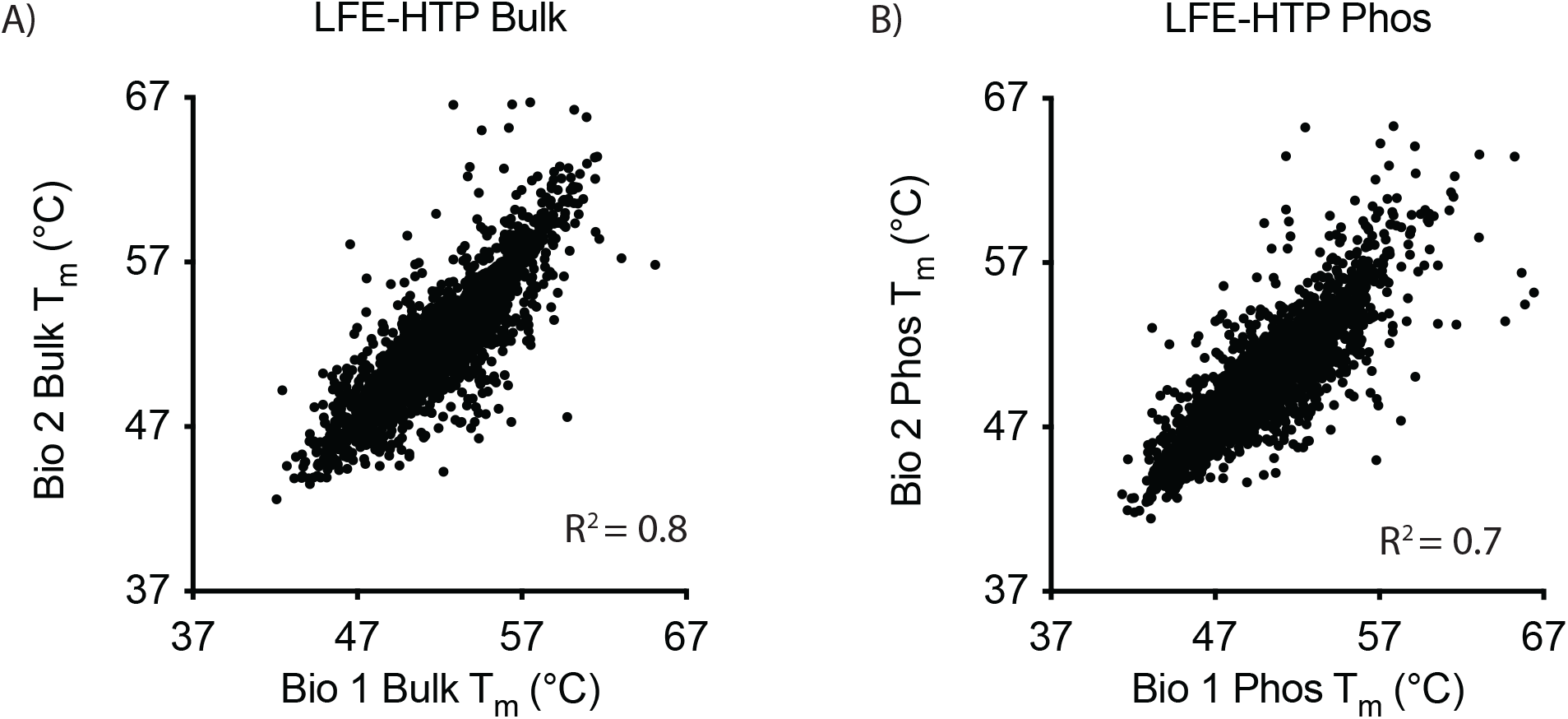
Representative T_m_ correlation plots from LFE-HTP analysis of bulk, unmodified (**A**) and phosphoproteome (**B**).

**Suppl. Fig. 4.**
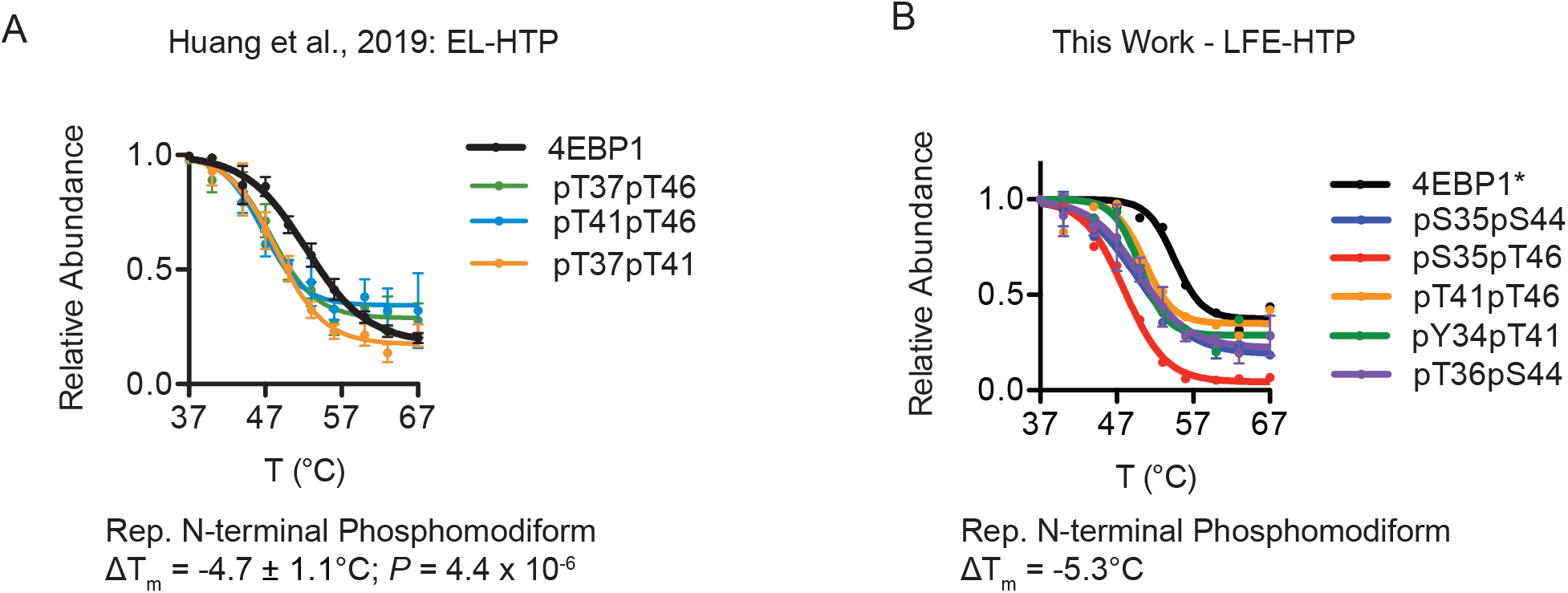
Comparison of 4EBP1 phosphomodiform melting profiles from Huang et al 2019 (**A**) and LFE-HTP (**B**). Only curve-fit with *R*^2^ > 0.8 are plotted, including cases where only one curve for specific bulk or phosphosites were detected and passed these criteria (denoted by an*). Curves and error bars correspond to mean and s.e.m.; ΔT_m_ *P*-values calculated with two-sided *t*-test.

**Suppl. Fig. 5:**
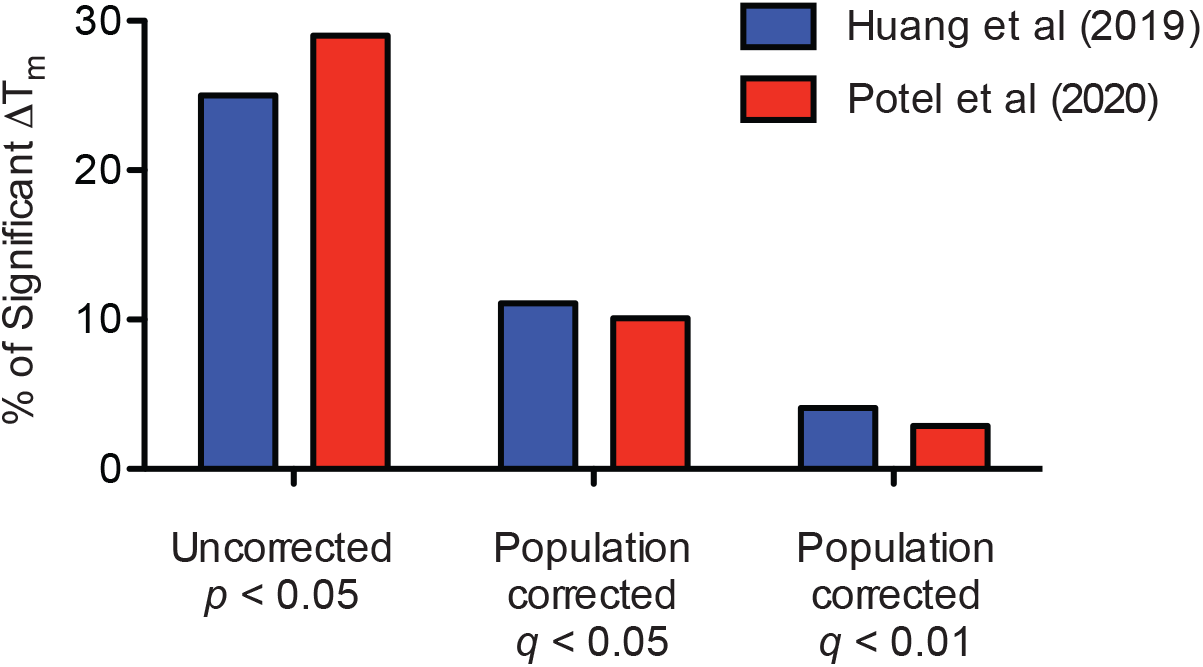
Comparing the percentages of significant ΔT_m_ values based on application of different statistical cutoffs for our original published dataset (blue) and the Potel et al dataset (red). The percentages of significant ΔT_m_ values with *p*-value < 0.05 are 25% (Huang) and 29% (Potel) of the two datasets using our published analysis pipeline. When we apply a population-level, multiple testing correction to generate *q*-values, the percentages of ‘significant’ ΔT_m_ values are reduced. Specifically, application of *q*-value < 0.05 produces 11% (Huang) and 10% (Potel) and *q*-value < 0.01 produces 4.1% (Huang) and 2.9% (Potel) significantly shifted ΔT_m_ values.

**Supplementary Table 1:** Bulk T_m_ values from LFE-HTP

**Supplementary Table 2:** Phosphosite T_m_ values from LFE-HTP

## Methods

### Cell Culture

HEK293T cells were propagated in DMEM with sodium pyruvate and L-glutamine (Corning) supplemented with 10% fetal bovine serum (R&D systems) and 1% penicillin-streptomycin (Thermo Fisher Scientific, GIBCO) and grown at 37°C in a 5% CO_2,_ humidified incubator.

### LFE-HTP Cell Harvesting and Proteomic Sample Preparation

For LFE (Label-Fractionate-Enrich) HTP samples, cells were lifted using TrypLE Express (Thermo Fisher Scientific - GIBCO) and neutralized following 5-minute incubation using complete media (DMEM + 10% FBS penicillin/streptomycin) and centrifuged at 1100 r.p.m. for 4 minutes. The cell pellet was reconstituted in 10 mL PBS containing protease and phosphatase inhibitors (Roche) and centrifuged again at 1100 RPM for 4 minutes. Following centrifugation, the cell pellet was resuspended in 1 mL PBS with inhibitors and distributed into thin-wall PCR tubes at 100 μL of cell suspension in each tube. Thermal denaturation was performed as previously described^3^, and the resulting cellular suspension was transferred to clean 1.5 mL microcentrifuge tubes and PCR tubes were additionally rinsed with 30 μL of PBS with inhibitors to ensure complete transfer of cellular suspension. Cellular suspension was next snap frozen in liquid nitrogen for 1 minute followed by thawing and re-equilibration back to room temperature. This freeze-thaw cycle was repeated 2 additional times and the soluble fraction of each lysate was generated by centrifugation at 21,130 *x g* for 30 minutes at 4°C. Supernatants were transferred to clean 1.5 mL microcentrifuge tubes, and protein was quantified in the supernatant for temperatures 37°C and 41°C by micro BCA assay (Thermo Fisher Scientific - Pierce). Following quantification, the average of the two lowest temperatures was taken and the volume equivalent to 30 μg of protein in the lowest temperature was moved from each temperature fraction into a clean 1.5 mL tube. Following distribution of protein, each tube was brought to a final volume of 300 μL by addition of PBS with inhibitors, followed by precipitation with trichloroacetic acid (TCA) (Sigma) to a final concentration of 25%, vigorously vortexed and incubated on ice overnight. TCA precipitates were centrifuged at 21,130 *x g* for 30 minutes at 4°C, washed twice in 500 µL of ice-cold acetone, and centrifuged at 21,130 *x g* for 10 minutes after each wash. Following precipitation and washes, pellets were allowed to completely dry at room temperature. Dry pellets were re-suspended in 100 μL of 100 mM TEAB, 0.5% SDS and reduced with 9.5 mM tris-carboxyethyl phosphine (TCEP) for 60 minutes at 55°C. Following reduction of disulfide bonds with TCEP, the denatured protein mix was centrifuged at 21,130 *x g* for 5 minutes then alkylated with 4.5 mM iodoacetamide (IAA) for 30 minutes in the dark at room temperature. After reduction and alkylation of disulfide bonds, the denatured protein mixture was precipitated out of solution by addition of 600 µL of ice-cold acetone and placed in the -20°C freezer overnight. The following day precipitated proteins were centrifuged at 8,000 *x g* for 10 minutes to pellet precipitated protein. Following centrifugation supernatant was decanted off and pellets were allowed to air-dry at room temperature. Once dry, protein pellets were reconstituted in 100 µL 100 mM TEAB and CaCl_2_ was supplemented to a final concentration of 1 mM, 1 µg of sequencing grade Trypsin (Promega) was added, and reactions were placed in the dark on a thermal mixer (Eppendorf) set to 37°C and shaking at 850 r.p.m. for 16 hours. The next day, digested samples were centrifuged at 21,130 *x g* for 10 minutes and proceeded to TMT labeling of digested samples.

### LFE-HTP TMT Labeling, Fractionation, and Phosphopeptide Enrichment

For LFE-HTP (Label-Fractionate-Enrich) experiments, TMT labeling was performed generally as per manufacturer’s protocol. Briefly, each TMT tag was re-suspended in 164 μL anhydrous acetonitrile with intermittent vortexing for 10 minutes. Following resuspension, 41 μL was added to corresponding temperatures (TMT-126 = 37°C; four separate aliquots of each temperature for subsequent desalting and fractionation) and labeling reaction was allowed to proceed for 1 hour at room temperature. Reactions were quenched by addition of 8 μL of 5% hydroxylamine in 100 mM TEAB and incubated for 15 minutes. Labeled temperature fractions were pooled, desalted on 1cc/50 mg C18 SepPAK columns (Waters # WAT054955) on a vacuum manifold and desalted peptides were dried down in a speedvac. Dried peptides were reconstituted in 300 µL of 0.1% TFA in H_2_O, high-pH reverse phase spin-columns (Thermo fisher scientific - Pierce) were equilibrated, and samples fractionated per manufacturer’s instructions into 8 fractions, 2 washes and a flow-through fraction (11 total). Separate samples from the same fractions were then combined and dried. Peptide fractions were reconstituted in 200 µL of 5% acetonitrile, 0.1% TFA in water, and 10 µL was removed for bulk HTP analysis. The remaining fractionated labeled peptides dried and re-dissolved in 40% acetonitrile, 6% TFA in water before phosphopeptide enrichment with Titansphere 5 µm TiO_2_ beads (GL Sciences). Titansphere TiO2 beads (GL Sciences) were reconstituted in buffer containing 80% acetonitrile, 6% TFA, and 2,5-dihydroxybenzoic acid (20 mg/mL) and rotated for 15 min at 25°C. Equal amount of beads slurry (∼5:1 beads-to-peptide ratio based on concentration of peptides in 37°C aliquot) was added to each temperature aliquot of reconstituted peptides and rotated for 20 mins 25°C. Beads were then washed twice with higher percentage of acetonitrile (10% and 40%) in 6% TFA and supernatant was removed by centrifugation at 500 *x g* for 2 min. Washed beads were then added to self-packed stage tip with C8 SPE (Sigma Aldrich) and washed once more with 60% acetonitrile in 6% TFA. Phosphopeptides were first eluted with 5% NH_4_OH, then 10% NH_4_OH, 25% acetonitrile, and dried with speedvac. Dried phosphopeptides were reconstituted in 5% acetonitrile, 1% TFA, desalted with self-packed stage tip with C18 SPE (Sigma Aldrich), and dried with speedvac once more. The final processed phosphopeptides were reconstituted in 5% acetonitrile, 0.1% TFA in water for LC-MS^3^ analysis.

### LC-MS^3^ Analysis and Data Acquisition

High-pH reverse-phase fractions were run on a 4-hour instrument method with an effective linear gradient of 180 minutes from 5% to 25% mobile phase B with the following mobile phases: A: 0.1% formic acid in H_2_O, B: 80% acetonitrile/0.1% formic acid in water on a 50 cm Acclaim PepMap RSLC C18 column (Thermo Fisher Scientific #164942) operated by a Dionex ultimate 3000 RSLC nano pump with column heating at 50°C connected to an Orbitrap Fusion Lumos. Briefly, the instrument method was a data-dependent analysis and cycle time set to 3 seconds, total. Each cycle consisted of one full-scan mass spectrum (400-1500 m/z) at a resolution of 120,000, RF Lens: 60%, maximum injection time of 100 ms followed by data-dependent MS/MS spectra with precursor selection determined by the following parameters: AGC Target of 4.0e5, maximum injection time of 100 ms, monoisotopic peak determination: peptide, charge state inclusion: 2-7, dynamic exclusion 10 sec with an intensity threshold filter: 5.0e3. Data-dependent MS/MS spectra were generated by isolating in the quadrupole with an isolation window of 0.4 *m*/z with CID activation and corresponding collision energy of 35%, CID activation time of 10 ms, activation Q of 0.25, detector type Ion Trap in Turbo mode, AGC target of 1.0e4 and maximum injection time of 120 ms. Data-dependent multi-notch MS^3^ was done in synchronous precursor selection mode (SPS, multi-notch MS^3^) with the following settings: Precursor selection Range; Mass Range 400-1200, Precursor Ion Exclusion Properties *m*/z Low: 18 High: 5, Isobaric Tag Loss Exclusion Properties: TMT. Number of SPS precursors was set to 10 and data-dependent MS^3^ was detected in the Orbitrap (60,000 resolution, scan range 120-500) with an isolation window of 2 *m*/z HCD activation type with collision energy of 55%, AGC target of 1.2e5 and a maximum injection time of 150 ms. Raw files were parsed into MS1, MS2 and MS3 spectra using RawConverter.

### Data Analysis

Data generated were searched using the ProLuCID algorithm in the Integrated Proteomics Pipeline (IP2) software platform. Human proteome data were searched using a concatenated target/decoy UniProt database. Basic searches were performed with the following search parameters: HCD fragmentation method; monoisotopic precursor ions; high resolution mode (3 isotopic peaks); precursor mass range 600-6,000 and initial fragment tolerance at 600 p.p.m.; enzyme cleavage specificity at C-terminal lysine and arginine residues with 3 missed cleavage sites permitted; static modification of +57.02146 on cysteine (carboxyamidomethylation),

+229.1629 on N-terminal and lysine for TMT-10-plex tag; 4 total differential modification sites per peptide, including oxidized methionine (+15.9949), and phosphorylation (+79.9663) on serine, threonine, and tyrosine (only for phospho-enriched samples); primary scoring type by XCorr and secondary by Zscore; minimum peptide length of six residues with a candidate peptide threshold of 500. A minimum of one peptide per protein and half-tryptic peptide specificity were required. Non-unique peptides were included in search. Starting statistics were performed with a Δmass cutoff = 10 p.p.m. with modstat, and trypstat settings. False-discovery rates of peptide (sfp) were set to 1%. TMT quantification was performed using the isobaric labeling 10-plex labeling algorithm, with a mass tolerance of 5.0 p.p.m. or less. Reporter ions 126.127726, 127.124761, 127.131081, 128.128116, 128.134436, 129.131417, 129.13779, 130.134825, 130.141145, and 131.13838 were used for relative quantification.

### Statistical and Melting Curve Analysis

The LFE-HTP dataset is comprised of n = 4 independent experiments derived from materials from independent cell cultures. Each independent experiment corresponded to n = 22 unique mass spectrometry measurements (n = 11 for fractionated bulk proteome, n = 11 for fractionated phospho-enriched proteome). For melting curve analysis, all detected phosphopeptides that map to the same phosphorylation sites were grouped together and given a new identifier (Gene_pSite). For example, 4 different tryptic peptides mapping to the same GAPDH phosphorylation site pS210 were often detected: R.DGRGALQNIIPAS(79.9963)TGAAKAVGK.V, R.DGRGALQNIIPAS(79.9963)TGAAK.A, R.GALQNIIPAS(79.9963)TGAAKAVGK.V, and R.GALQNIIPAS(79.9963)TGAAK.A. These peptides were systematically re-labeled with the new custom identifier “GAPDH_pS210”. TMT reporter ion intensities of phosphopeptides with the same identifier were combined for each temperature fraction from the same MS run. Fold change values were calculated using the lowest temperature fraction as the reference. Similarly, for the bulk unmodified proteome, TMT reporter ion intensities of all unmodified peptides mapped to the same protein were combined and fold-change values relative to the lowest temperature condition were calculated. To generate unmodified protein and phosphosite-specific melting curves, relative fold-changes as a function of temperature was fitted to the equation derived from the chemical denaturation theory using R according to previously described^1^. T_m_ values were calculated at which the sigmoidal curve crosses the 0.5 fold-change level. Only T_m_ values calculated from melting curves with curve R^2^ > 0.8 were used in subsequent analyses. Shift in T_m_ values (e.g. ΔT_m_) induced by phosphorylation were determined by subtracting the T_m_ of the bulk, unmodified protein from the T_m_ of the phosphomodiform:

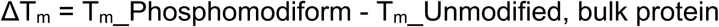

ΔT_m_s could only be calculated for phosphosites belonging to proteins that were also detected in the unmodified proteome. T_m_ values calculated by the R script of all unmodified proteins and phosphorylation sites are summarized in Supplementary Tables 1-2. For all proteins highlighted, normalized relative abundance ratios were plotted and fitted with variable-slope (four-parameter) curve fit function in Prism 9 (GraphPad). For all proteins and phosphomodiforms highlighted in which the R script was unable to generate a curve fit, the variable-slope (four-parameter) curve fit function Prism 9 was used to generate a T_m_ value for ΔT_m_ calculation. To calculate p-value for ΔT_m_ values, two-sided student’s t-test was used to compare the mean T_m_ ± s.e.m. for a defined phosphomodiform of interest to the mean T_m_ ± s.e.m. for the bulk, unmodified protein pool, both derived from independent measurements from unique MS runs.

## Notes

### Competing Interest Statement

The authors have declared no competing interest.

